# Oxytocin effects on the resting-state mentalizing brain network

**DOI:** 10.1101/465658

**Authors:** Haiyan Wu, Chunliang Feng, Xiaping Lu, Xun Liu, Quanying Liu

## Abstract

Oxytocin(OT) has effects in both human behavior and in the brain, which is not limited in the specific brain area but also with the potential effect on connectivity with other brain regions. Evidence indicate that the effects of OT on human behavior are multifaceted, such as trust behavior, decrease anxiety, empathy and bonding behavior. Since the vital role of mentalizing in understanding others, here we proposed and tested that whether OT has a general effect on theory of mind brain network which is associated to the effect of related social behavioral and personality traits. Used a randomized, double-blind placebo-controlled group design, we investigated the resting-state functional magnetic resonance imaging after intranasal OT or placebo. The functional connectivity (FC) maps with seed in left temporoparietal junction (lTPJ) and right TPJ showed that OT significantly increased connectivity between rTPJ and default attention network (DAN), while decreased the FC between lTPJ and medial prefrontal network (MPN). With implementing machine learning approach, we further reported satisfactory classification accuracy that Identified altered FCs of TPJ can classify OT and PL group. Moreover, individual’s empathy trait can modulate the FC between left TPJ and right RECT, which was positively correlated with empathic concern in PL group whereas lTPJ-rRECT negatively correlated in OT group. These results demonstrate that OT has significant effect on FC with lTPJ and rTPJ, brain regions critical for mentalizing, and the empathy concern can modulate the FC. These findings add to our understanding of the neural mechanisms by which OT modulates social behaviors, especially in social interaction involving mentalizing.

## INTRODUCTION

Oxytocin (OT) is a neuropeptide which has been shown to modulate complex emotional and social behaviors (Bartz et al., 2011), and mediate the relevant neural activity (Kirsch et al., 2005) and functional connectivity (Bethlehem et al., 2013; Dodhia et al., 2014). It has been evident that oxytocin modulates various social emotions and behaviors (Alvares et al., 2010; Hu et al., 2015; Kosfeld et al., 2005; Labuschagne et al., 2010a; Riem et al., 2011; Scheele et al., 2013; Shamay-Tsoory et al., 2009; Striepens et al., 2012; Theodoridou et al., 2009). For example, behavioral studies have shown that oxytocin administration induces more trust behaviors (Kosfeld et al., 2005), and decrease social distance from others (Preckel et al., 2014), and even more prosocial behaviors to social rejected others (Riem et al., 2013a). Neuroimaging studies have convergently shown oxytocin modulates brain activity in social behaviors (Wittfoth-Schardt et al., 2012), such as emotion recognition (Hurlemann et al., 2010), social memory (Rimmele et al., 2009) as well as prosocial behaviors (Mikolajczak et al., 2010). For instance, OT administration changes the brain activity in response to infant crying, including decreased the activity in amygdala and increased activity in both insula and inferior frontal gyrus (Riem et al., 2011), which may be attributed to increasing empathic responses. Similarly, OT-administrated subjects showed a reduced response of the right amygdala to angry, happy and fearful faces (Domes et al., 2007a). Accordingly, efforts have also been dedicated to its therapeutic potential in different kinds of social dysfunction disorders patients, in both behavioral and neural levels (Anagnostou et al., 2012; Labuschagne et al., 2010b; Modi and Young, 2012; Neumann and Landgraf, 2012). These results suggest that OT effect is specifically germane to social stimulus, and to the brain network activated by the stimulus.

While such attempts aim to understand the neurobiological basis of OT effect on social cognition through task-related functionally brain activities, there are growing interests in understanding brain network changes in task-free state, which are far less thoroughly investigated. To date, some studies in humans have investigated the effects of oxytocin on resting-state functional magnetic resonance imaging (rsfMRI). This body of work mainly motivated by hypothesis about the OT effect on emotion and amygdala pathway. For example, individuals with oxytocin misuse are with stronger connectivity between right amygdala and dorsal anterior cingulate cortex, relative to the control group (Kovacs and Keri, 2015). With a closer examination of amygdala subregions, Eckstein *et al* (2017) found OT exerts different amygdala subregion-specific connectivity patterns in resting state. In social anxiety disorder patients, oxytocin enhanced amygdala and rostral anterior cingulate cortex (ACC)/medial prefrontal cortex (mPFC) connectivity (Phan, 2014). Another related study also showed similar OT effect in healthy participants such that oxytocin significantly increased connectivity between amygdalae and rostral medial frontal cortex (Sripada et al., 2013). Of these studies investigating the effects of oxytocin on resting-state fMRI and connectivity, amygdala related brain network has been qualified to be affected by OT.

Although it has been reported that human oxytocin could modulate both social cognition and resting state brain connectivity, how oxytocin modulates other social brain networks engaged in these social behaviors (e.g., social inference, empathy and reward system), is still largely unclear. Notably, few study has focused on relationship between OT administration and mind-reading related brain network changes. Mind-reading is a process in which people draw internal states inference and the weight those inference with context information from external states (e.g. expressions, gestures, signals, etc.) (Realo et al., 2003). Large amounts of studies have shown that OT administration can improve mind-reading (Domes et al., 2007b; Radke and de Bruijn, 2015; Riem et al., 2014b; Voorthuis et al., 2014). For example, using the Reading the Mind in the Eyes Test (RMET), the accuracy was significantly improved after oxytocin administration (Domes et al., 2007c). One key cortical brain region involved in mind-reading functions is temporoparietal junction (TPJ) (Saxe and Wexler, 2005). Causal evidence showed that brain damage in this region could lead to dysfunctions in both moral decisions (Lamm et al., 2007) and empathy tasks (Benedetti et al., 2009; Cheon et al., 2011; Decety et al., 2008a; Dodell-Feder et al., 2011; Jackson et al., 2006; Lamm et al., 2011). Further evidence or meta-analysis showed separately functions in TPJ (Krall et al., 2015a; Kubit and Jack, 2013; Lombardo et al., 2011; Mitchell, 2008), however, another experimental design study showed support to the theory of mind hypothesis in TPJ (Young et al., 2010). Further, that TPJ connected brain has also been modulated by both empathy and social behaviors (Emonds et al., 2014). Clearly, most of the aforementioned studies show the connections between theory of mind, empathy and TPJ.

In light of those OT effect in mind reading, we speculate that the brain connectivity by which people perceive (perspective taking tasks) and infer about other’s mind (theory of mind tasks) could be modulated by OT. Since previous neuroimaging studies have shown individual difference in OT effect on brain activity and its connectivity (Eckstein et al., 2017; Riem et al., 2014a; Riem et al., 2013b), we would also take into account of individual difference, such as empathy trait.

The Interpersonal Reactivity Index (IRI) (Davis, 1980; Davis, 1983) is a widely used scale to measure dispositional empathy. It has four subscales (i.e., perspective taking, fantasy, empathy concern, and personal distress). The perspective-taking scale assesses spontaneous attempts to adopt the perspectives of other people and see things from their point of view. Items on the fantasy scale measure the tendency to identify with characters in movies, novels, plays and other fictional situations. The other two subscales explicitly tap respondents’ chronic emotional reactions to the negative experiences of others. It has been suggested that the mind-reading accuracy was increased after oxytocin administration for those lower in empathic concern scale of the IRI (Radke and de Bruijn, 2015). We thus measure the empathic trait of individual participant to tap into the possible moderation effect of empathetic trait on social brain network in terms of the resting state functional connectivity.

The aim of this study was to explore the effects of oxytocin on TPJ related functional connectivity in resting state. With the increasing concern about the individual difference and context effect in oxytocin effect on social cognition (Bartz et al., 2011; Churchland and Winkielman, 2012; Olff et al., 2013), we believe that the OT administration could lead to both broad influence of general state change and individual difference on resting brain network. Given prior evidence that oxytocin enhances mind reading accuracy and social salience hypothesis of oxytocin (Shamay-Tsoory and Abu-Akel, 2016), we hypothesized that OT administrated group would exhibit increased connectivity between TPJ to other social brain regions, relative to the group under placebo, and show moderation effect of individual personality traits.

## METHODS

### Participants

The study’s participants included 59 right-handed male college students (age range 19~26 years, and education range 13~18 years), who were recruited via an online recruiting system. All participants filled out a screening form, and participants were included in the study only if they confirmed they were not suffering from any significant medical or psychiatric illness, not using medication, not drinking and/or smoking in a daily basis. Participants were instructed to refrain from smoking or drinking (except water) for 2 hours before the experiment. All experiments were conducted in Beijing Normal University. Participants received full debriefing on completion of the experiment. All participants provided written consent, and the study protocol was approved by Institutional Review Board (IRB) of Beijing Normal University.

### Study Design and Drug Administration

We used a double-blind placebo-controlled group design to investigate the effects of a single dose of intranasal oxytocin on the resting-state functional connectivity. Participants randomly assigned to oxytocin (OT) group or the placebo (PL) group (Table 1). Each participant visited twice, with the first visit to fill in questionnaires (e.g., the Interpersonal Reactivity Index (IRI)) and the second visit for MRI scanning.

**Table 1.** Demographic and Clinical Characteristics.

|  | **OT group** | **PL group** | **t-score<sup>a</sup>** | **p-value** |
| --- | --- | --- | --- | --- |
| age (year) | 22.86 (1.57) | 22.79 (2.27) | 0.134 | 0.894 |
| body weight (kg) | 65.87 (9.16) | 68.21 (9.37) | -0.970 | 0.336 |
| IRI total scores | 93.83 (10.81) | 95.10 (9.09) | -0.478 | 0.634 |
| Perspective taking<sup>b</sup> | 22.79 (3.60) | 22.03 (2.91) | 0.883 | 0.381 |
| Fantasy<sup>b</sup> | 23.14 (5.83) | 25.17 (5.06) | -1.419 | 0.161 |
| Empathic concern<sup>b</sup> | 25.83 (4.54) | 26.21 (4.16) | -0.332 | 0.741 |
| Personal distress<sup>b</sup> | 22.07 (3.94) | 21.69 (4.40) | 0.346 | 0.731 |

Data are expressed as mean ( ± SD; standard deviation); n=30 for oxytocin group; n=29 for placebo group.
Abbreviations: OT, oxytocin; PL, placebo; IRI, Interpersonal Reactivity Index;
<sup>a</sup> t-test, two tailed.
<sup>b</sup> Data missing from one OT group (n=29).

Each participant arrived for the second visit 1 hour before MRI scanning. Participants did not consume food, caffeine or alcohol 1 hour before the MRI scanning. At 45 minutes before the MRI scanning, a single dose of 24 IU oxytocin or placebo was administered intranasally by three puffs of 8 IU per nostril. The puffs were delivered to each nostril in an alternating order and with a 45-second break between each puff to each nostril. The participants have 5-minute resting-state scan before any tasks. Each subject was positioned supine in the MRI scanner with his head in a comfortable restraint to reduce movement. Subjects were instructed to keep their eyes closed, relax and let their minds wander, but not to fall sleep. Following the resting-state scan, participants performed N different cognitive tasks. These task-relevant data were analyzed separately from the resting-state results.

### MRI Acquisition

A 3T Siemens Tim Trio scanner with 12-channel head coil was used to perform MRI. Functional images employed a gradient-echo echo-planar imaging (EPI) sequence that is sensitive to blood oxygenated level-dependent (BOLD) contrast (40 ms TE, 2 s TR, 90° flip, 210 mm FOV, 128 × 128 matrix, 25 contiguous 5 mm slices parallel to the hippocampus and interleaved). We also acquired from all participants whole-brain T1-weighed anatomical reference images (2.15 ms TE, 1.9 s TR, 9° flip, 256 mm FOV, 176 sagittal slices, 1 mm slice thickness, perpendicular to the anterior-posterior commissure line).

### Data preprocessing

fMRI data preprocessing was performed using Statistical Parametric Mapping software (SPM12: Wellcome Trust Centre for Neuroimaging, London, UK). The functional image time series were preproessed to compensate for slice-dependent time shifts, motion corrected, and linearly detrended, then coregistered to the anatomical image, spatial normalized to Montreal Neurological Institute (MNI) space (http://www.bic.mni.mcgill.ca/ServicesAtlases/HomePage) and spatially smoothed by convolution with an isotropic Gaussian kernel (FWHM=6 mm). The fMRI data were high-pass filtered with a cutoff of 0.01 Hz. The white matter (WM) signal, cerebrospinal fluid (CSF) signal and global signal, as well as the 6-dimensional head motion realignment parameters, the realignment parameters squared, their derivatives, and the squared of the derivatives were regressed. The resulting residuals were then low-pass filtered with a cutoff of 0.1 Hz.

### ICA-based RSN extraction

We used a data-driven approach to extract RSNs from both OT and PL groups, to investigate the effects of OT on large-scale functional networks. Specifically, 14 regular RSNs were extracted in each individual fMRI dataset by using ICA-based approach(Beckmann et al., 2009), combining a template matching strategy (Liu et al., 2017), including cingulo-opercular network (CON), default mode network (DMN), dorsal attention network (DAN), dorsal somatomotor network (DSN), auditory network (AN), language network (LN), left frontoparietal network (lFPN), right frontoparietal network (rFPN), medial prefrontal network (MPN) and lateral prefrontal network (LPN), ventral attention network (VAN), ventral somatomotor network (VSN), visual foveal network (VFN) and visual peripheral network (VPN). The group-level RSN maps for both groups were obtained by using a voxel-wise non-parametric permutation test by FSL (http://fsl.fmrib.ox.ac.uk/fsl/fslwiki) with 5000 permutations. Besides, two-sample unpaired t-test between OT and PL group was applied to investigate the topographic changes induced by OT.

### Seed-based functional connectivity analysis

The analysis pipeline was conducted in MATLAB and was similar to that employed in our previous research (Wu et al., 2015). We averaged the signals in left and right TPJ masks, which were defined by previous literature (Alcala-Lopez et al., 2017). The temporal correlations between seed region and the rest of the brain were examined, subject by subject. To generate the TPJ connectivity map for each group, one-sample t-tests were performed for both OT and PL group by running *randomise* function in FSL. To compare the FC between OT and PL group, two-sample unpaired t-test was applied. The resulting maps were thresholded at p<0.05 with threshold-free cluster enhancement (TFCE) correction (Smith and Nichols, 2009). The clustered regions with peak MNI coordinates were exported by *cluster* function in FSL. We used the FCs between TPJ and these identified regions for the following analyses.

### Classification of OT and PL with TPJ functional connectivity

To examine whether the distinction the TPJ connectivity patterns from OT and PL were consistent across participants, we used a linear support vector machine (SVM) classifier and the identified FC as input features to distinguish OT and PL. SVM identifies the weight of each FC that define a maximum margin hyperplane that best separates two classes of data (Dosenbach et al., 2010). We used leave-one-out cross-validation (LOOCV), where the classifier was trained on 58 participants (*svmtrain* function in MATLAB), then tested on the remaining one participant (svmclassify function in MATLAB). The performance was quantified using the accuracy rate, sensitivity and specificity based on the results of cross-validation (Zeng et al., 2012). The weight of each FC was recorded for all iterations of the cross-validation to examine the reliability of consensus functional connectivity discriminative power.

### Post-hoc correlation analyses

To investigate whether the participant’s dispositional empathy intermediate the oxytocin effect on TPJ connectivity, we calculated the partial correlation between each IRI subscale and FCs of the identified brain regions, by statistically removing the other subscales. The false discovery rate (FDR) correction were performed on p values.

## RESULTS

### Oxytocin effects on TPJ brain connectivity

We firstly obtained the FC maps with seed in left TPJ and right TPJ and examine the group-level FC maps in OT group and PL group, respectively (Figure 1). Clearly, we found that TPJ positively connected with the brain regions involved in default mode network (DMN) and medial prefrontal network (MPN), but negatively correlated with the brain regions in dorsal attention network (DAN). To quantify the oxytocin effects on the functional connectivity between TPJ and brain networks, we extracted 14 resting-state networks (RSNs) by ICA approach (supplementary Figure 1). Surprisingly, no topological differences of RSNs from OT and PL group were significant, which indicated that oxytocin does not change the topography of RSNs. We further calculate the FC between TPJ and 14 extracted RSNs, and proved that both left and right TPJ strongly synchronized with DMN and VAN, and anticorrelated with CON, DSN and AN (Figure 2A and 2B).

**Figure 1.**
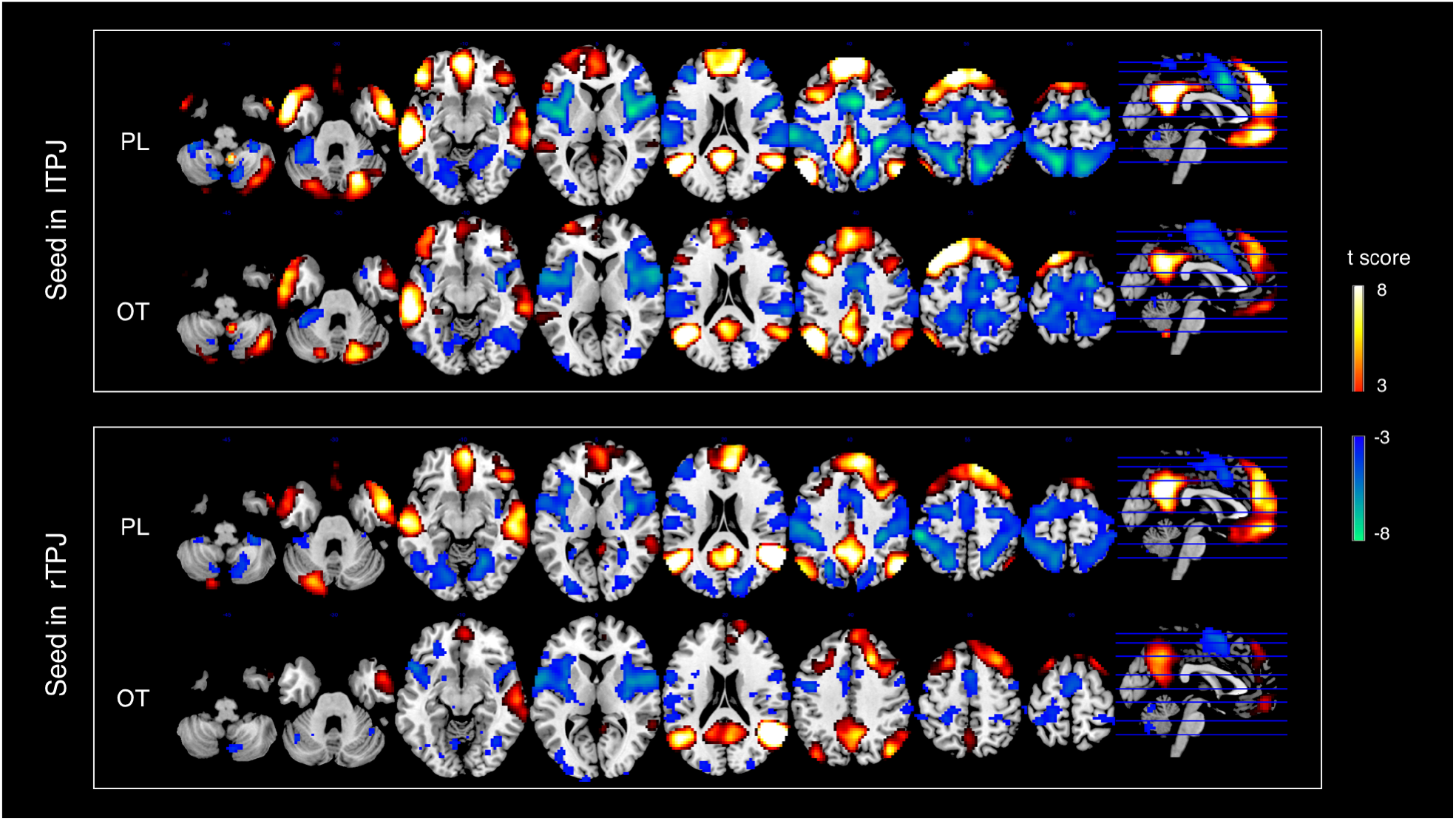
Functional connectivity with seed in left and right TPJ. Significant brain regions (p < 0.01, TFCE corrected) are indicated on the map. The brain regions labeled with yellow/red indicate positive correlation with seed region, while green/blue indicates negative correlation with seed region.

**Figure 2.**
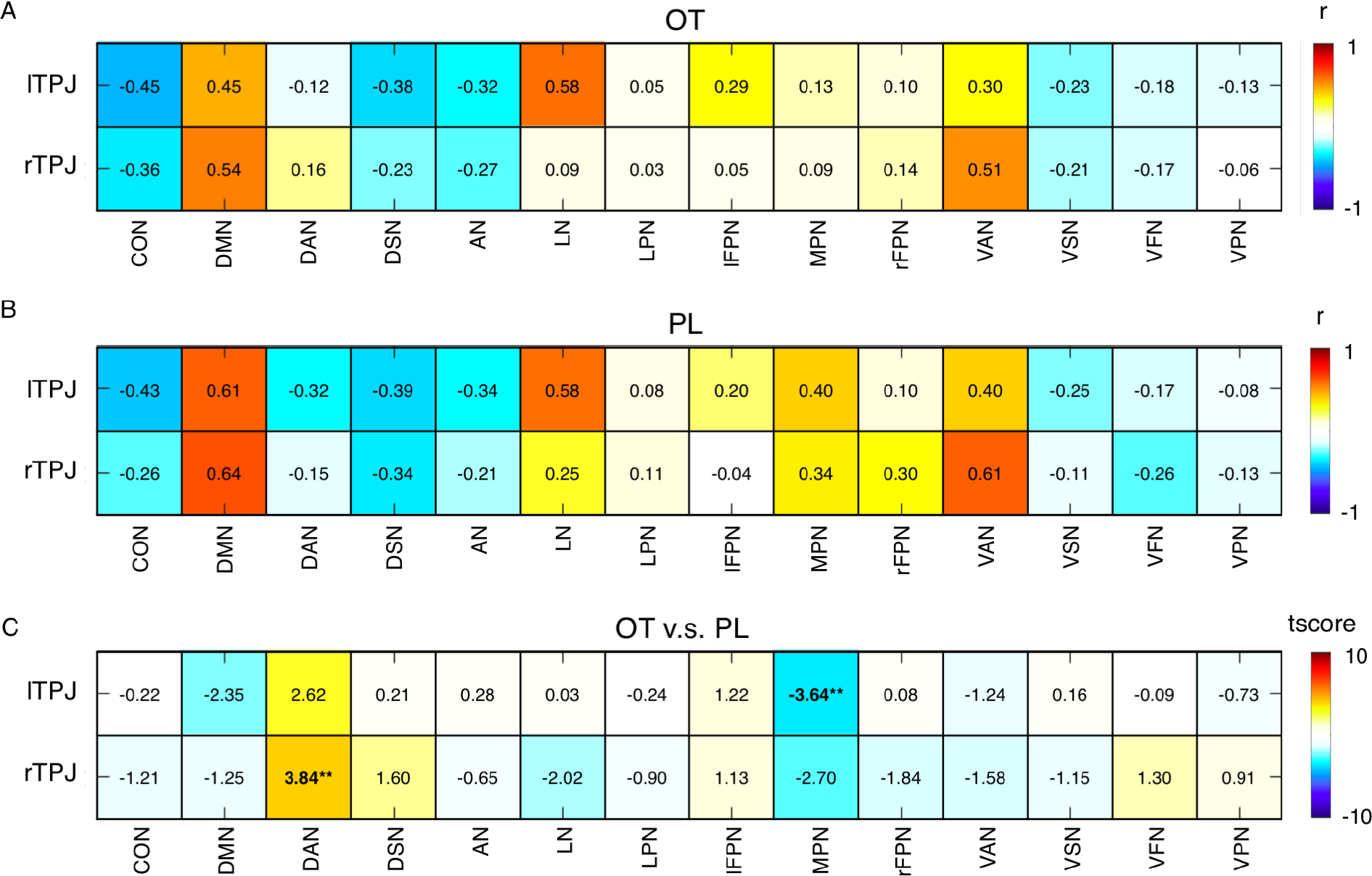
FC between TPJ and RSNs. A & B, the FC of left/right TPJ and 14 investigated RSNs for OT and PL group, respectively. The FC were calculated by Pearson correlation between averaged signals across these brain regions. C, t-scores from two sample t-test between TPJ-RSN connectivity. ** p<0.01. *Abbreviations*: CON, cingulo-opercular network; DMN, default mode network; DAN, dorsal attention network; DSN, dorsal somatomotor network; AN, auditory network; LN, language network, LPN, lateral prefrontal network; lFPN, left frontoparietal network; MPN, medial prefrontal network; rFPN, right frontoparietal network; VAN, ventral attention network; VSN, ventral somatomotor network; VFN, visual foveal network; VPN, visual peripheral network.

**Figure 3.**
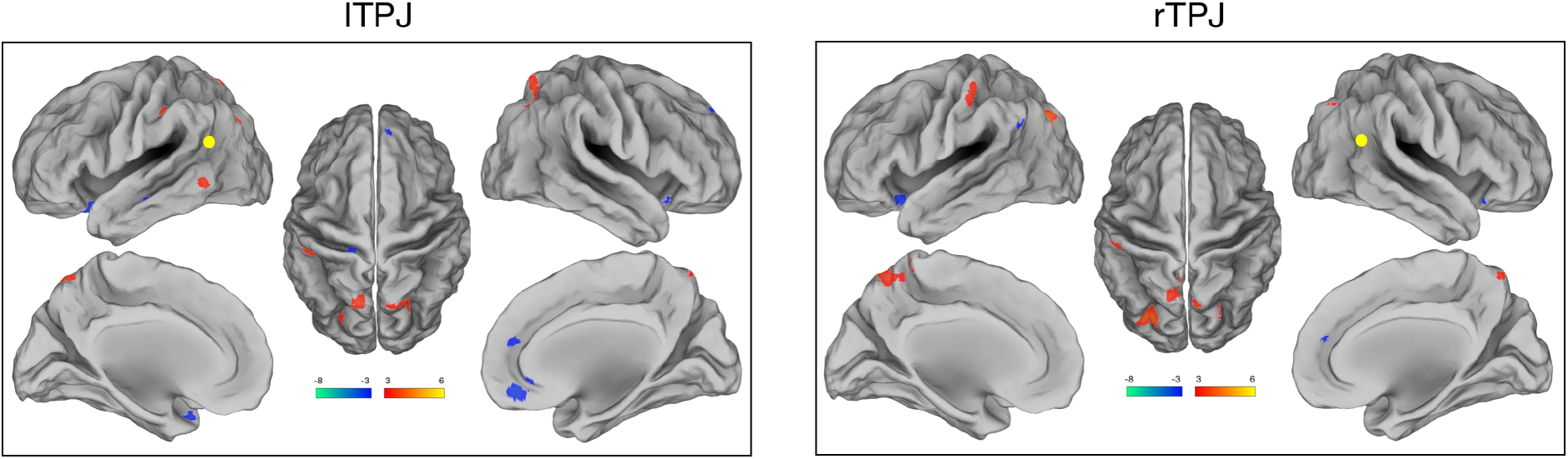
Comparisons between OT and PL group with seed in left and right TPJ, respectively. Seeds in lTPJ and rTPJ are depicted by the yellow dot. For visualization, the brain regions (|t-score|>3) are indicated on the map. The brain regions labeled with yellow/red indicate OT > PL, while green/blue indicates OT < PL. The peak MNI coordinate regions for each comparison are reported in Table 2. *Abbreviations*: TPJ, temporo-parietal junction; lTPJ, left temporo-parietal junction; rTPJ, right temporo-parietal junction; OT, oxytocin; PL, plaxebo; TFCE, threshold-free cluster enhancement.

Moreover, compared with PL group, the FC between lTPJ and MPN in OT is significantly weaker (t=-3.64, p=0.0083), whereas rTPJ-DAN in OT group is significant stronger (t=3.84, p=0.0044). All the reported p values were FDR corrected.

### Identified altered FCs

To compare the OT effect with placebo effect, we performed the between-group comparison (Figure 1). The resulting maps identified 8 brain regions (p<0.05, TFCE corrected; Table 1). We averaged the BOLD signals in these regions and then calculated the Pearson correlation between the identified brain regions and the corresponding seed region. Thus, we obtained 8 pairs of FCs with significant OT effects (ps<0.001, two-sample t-test), including lTPJ-lIPL (r=4.52), rTPJ-lPrec (r=5.36), rTPJ-rSPL (r=3.90), rTPJ-rPrec (r=3.82), rTPJ-lMOG (r=3.96) where the connectivity is higher in OT group, and lTPJ-rRECT (r=-4.35), rTPJ-lInsula (r=-4.11), rTPJ-lIFG (r=-4.12) where in intranasal OT reduced the coupling (Supplementary Figure 2).

### SVM classification based on the identified FCs

Using the identified eight FCs, the accuracy of classification was 74.58% with sensitivity of 73.33% and specificity of 75.86%. Moreover, the weights of input features were generated for each LOOCV training, which represents the contribution of each FC in classification, with a larger weight indicating larger contribution. Our results showed that lTPJ-rRect connectivity has the highest contribution. Intriguingly, the weights of these FCs were quite robust across subjects (see the mean and standard error of weights in Table 2), which indicates the reliable properties of TPJ related FC changes induced by OT administration.

**Table 2.**
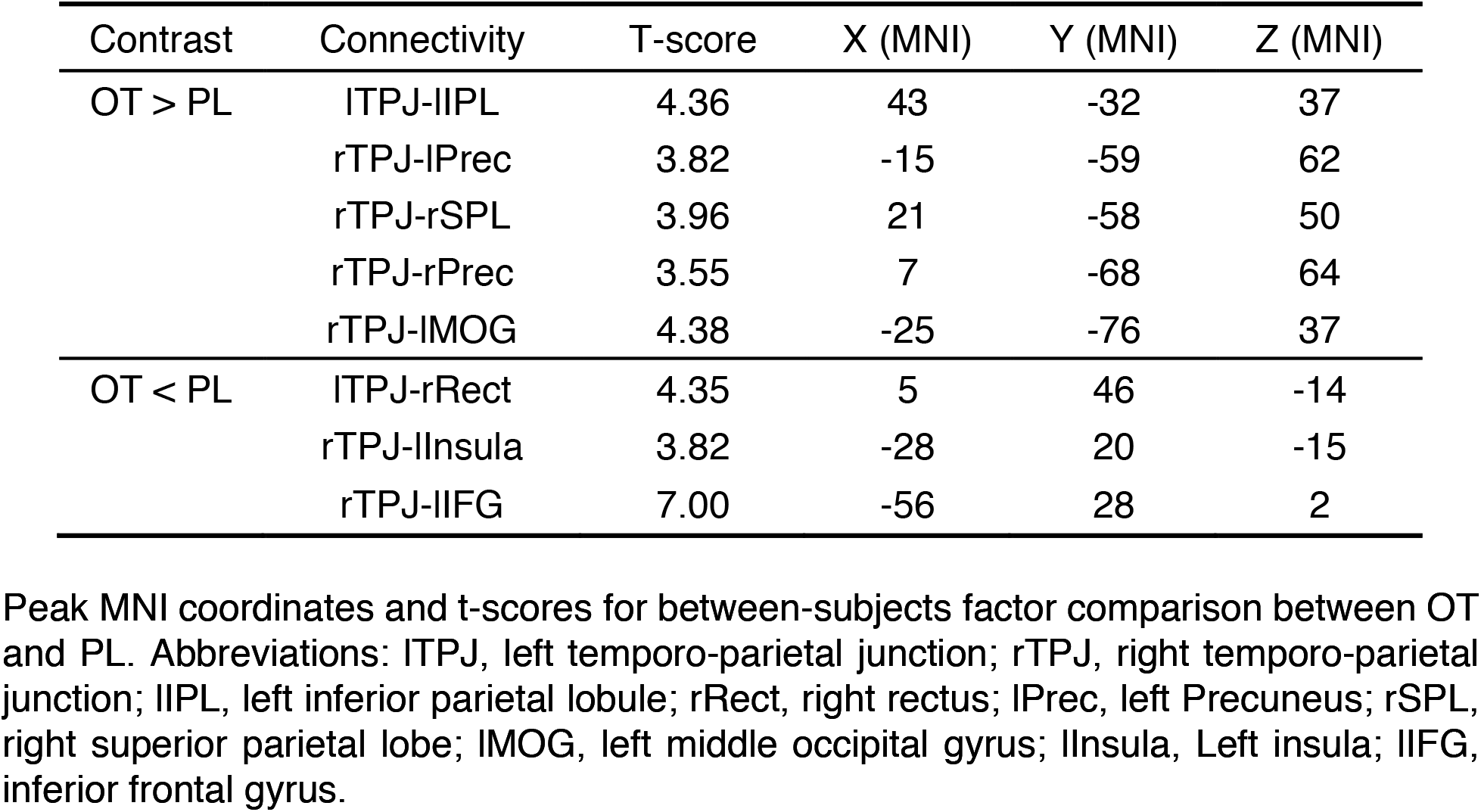
Results of between-subjects factor comparisons on the functional connectivity with TPJ4.

**Table 3.**
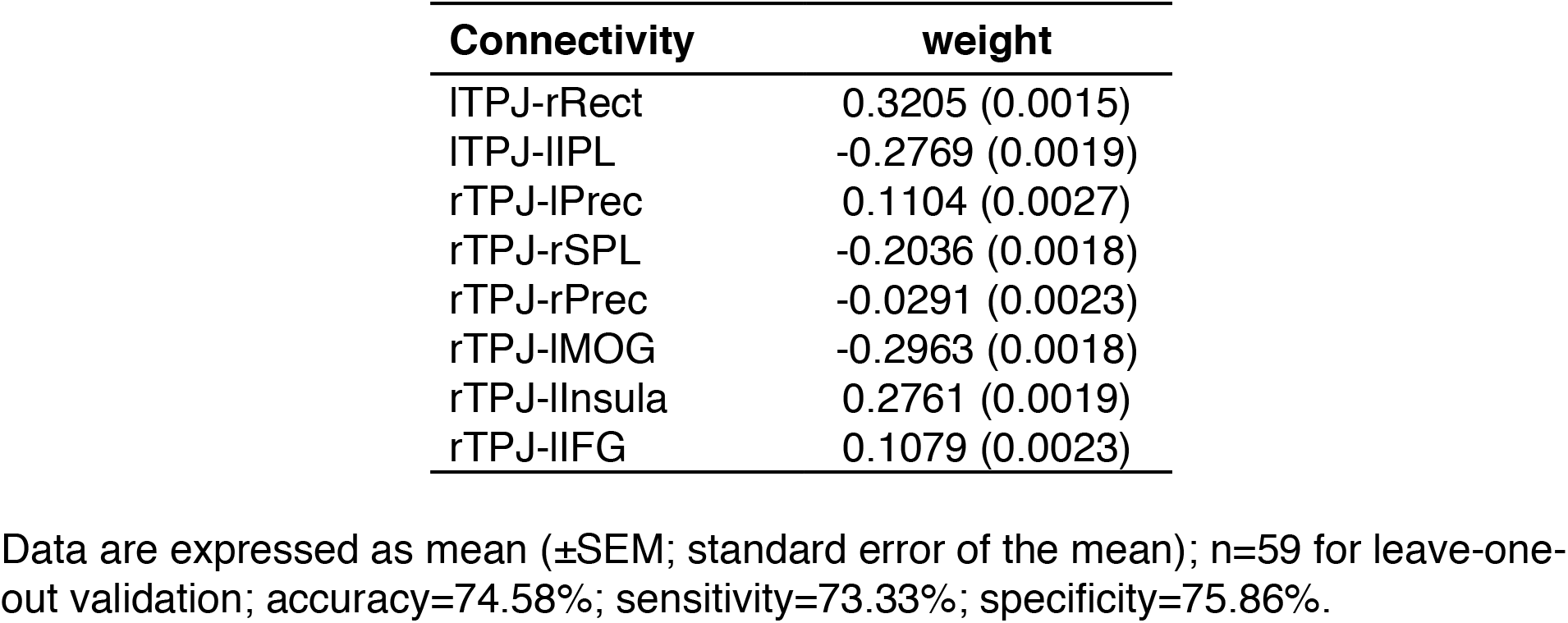
TPJ functional connectivity and the corresponding weight for SVM classification

### IRI intermediates the oxytocin effects on FC

To disentangle the influence of personal empathy on oxytocin effects, we calculated the partial correlation between each IRI subscale whilst controlling for the effect of other subscales. We found that the FC between left TPJ and right RECT positively correlated with empathic concern in PL group (r=0.47, p=0.059, Figure 4A), whereas lTPJ-rRECT negatively correlated in OT group (r=-0.51, p=0.029, Figure 4B). No FCs showed significant correlation with perspective taking, fantasy and personal distress.

**Figure 4.**
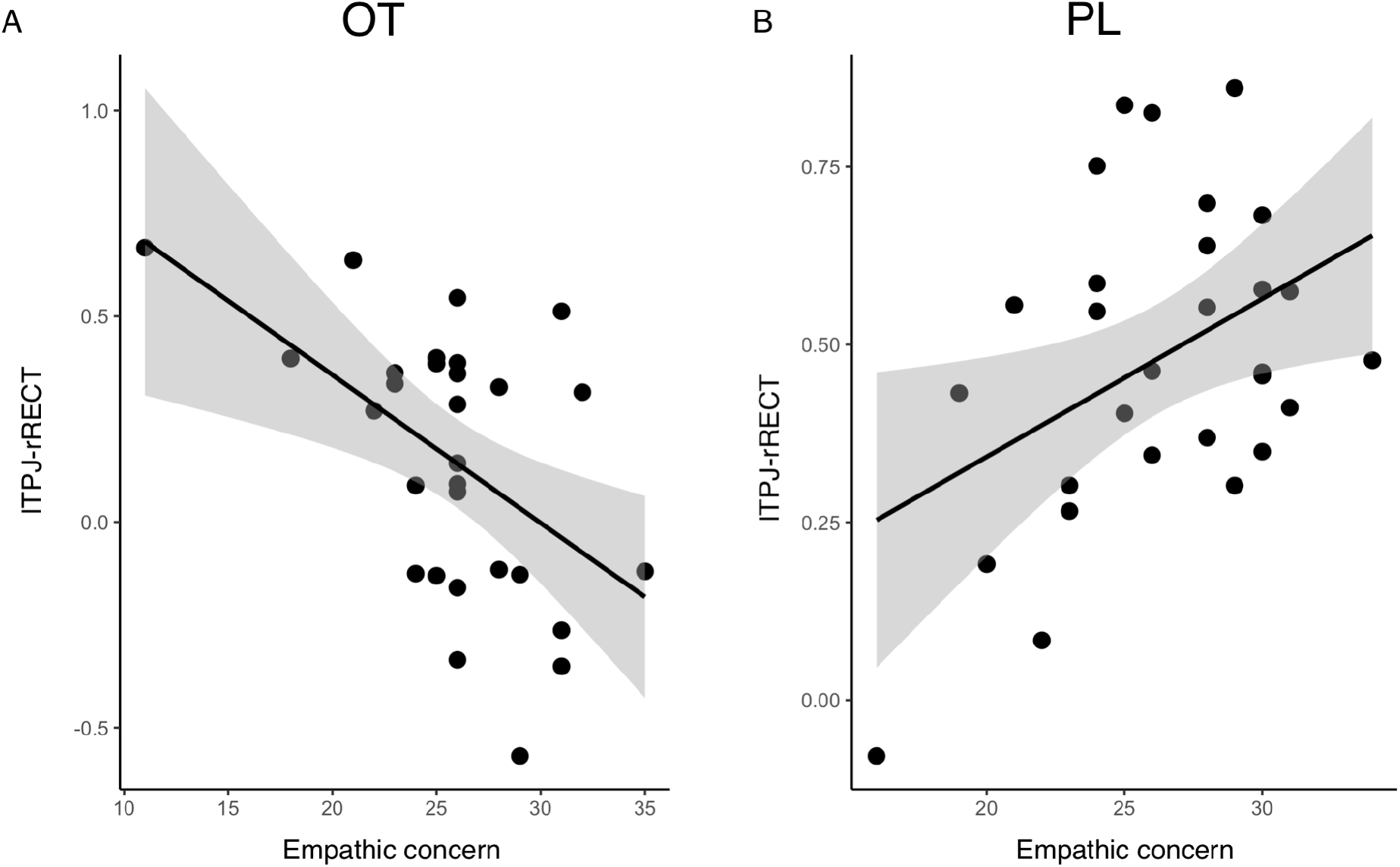
Empathic concern and TPJ-RECT functional connectivity. The scatter plot displays the linear correlation between the scores of empathic concern (x-axis) and the FC between left TPJ and RECT (y-axis), in OT group (left) and PL group (right). The shaded area represents the 95% confidence interval of the correlation line.

## DISCUSSION

Even playing a key role in social cognition, activity and connectivity of social brain networks in task-free state under oxytocin administration, are still not well understood. Recent studies have investigated amygdala - related brain network after OT administration. However, there are still other functions and brain networks relevant to social cognition that could give rise to these effects, which were not well investigated in previous work. In order to assess the OT effect on properties of mentalizing related brain networks, we examined OT effects on TPJ brain connectivity on putative brain networks and on social brain networks. The OT induced brain connectivity in social brain networks can be classified by distinguished FCs, which is modulated by individual traits.

### Interactions between TPJ and RSNs

Our results suggest that TPJ is generally positively correlated with DMN and MPN, but negatively correlated with DAN, independent of drug administration. The DMN is supposed to be the brain areas that are typically more active during rest state than during active task performance. However, it has been observed that DMN is partly overlapping with the social brain (Mars et al., 2012; Schilbach et al., 2008; Spreng et al., 2009; Washington et al., 2014; Xie et al., 2016). Our findings add further evidence on it based on the overlap between ICA-identified DMN and MPN (Figure 1). In line with the concept that TPJ connection with MPFC is vital in the understanding of other’s mental states (Li et al., 2014), we believe the observed generic positive connection between TPJ and MPN is related to monitor both self and other mental states. The negative connection between TPJ and DAN we observed, appears to echo existing findings with a dissociated effect on the DAN and ventral attention network (VAN) (Fox et al., 2006). It has been shown that the DAN is engaged in top-down orienting, while VAN activates for bottom-up salience detection (Corbetta and Shulman, 2002). As the TPJ acts as a part of VAN (Corbetta et al., 2000), the negative correlation reflects spontaneous neuronal activity between the two attention system (VAN and DAN) are relatively deviated (Fox et al., 2006). For example, previous work has shown decreased TPJ activity when cognitive demands are imposed and the DAN is engaged (Shulman et al., 2003).

### OT effects on TPJ functional connectivity

It is interesting that oxytocin does not change the ICA-identified topography of RSNs, but changes the FCs between TPJ and RSNs. Specifically, the PL group had stronger left TPJ and MPN connection, whereas OT group showed stronger connection between right TPJ and DAN. In accordance with our initial hypotheses that TPJ connection may play an essential role in self-other mental state monitoring process, the weaker connection in the PL group may due to the less sensitivity of social signal. Conversely, the stronger connection between right TPJ and DAN in OT group may be related to the enforced social attention, which provide the support for social salience hypothesis of oxytocin. Social inference and attention functions are considered as two aspects of function in TPJ (Krall et al., 2015b), where is also brain area that attention and memory information converged (Carter and Huettel, 2013). Since attention or working memory studies showing causality activity between VAN and DAN (Sridharan et al., 2007) and the essential role of right TPJ in theory of mind (Hooker et al., 2010; Young et al., 2010), the enhancing rTPJ and DAN connection might be explained by increased social perception function.

Importantly, it seems that the significant FC differences between PL and OT group provides further evidence of the social attention hypothesis. In particular, right TPJ connection with nearby brain regions such as precuneus, inferior parietal lobule, and superior parietal lobule, is stronger in OT group. These brain regions are proposed to play critical role in salient detection or attention sustaining (Singh-Curry and Husain, 2009). Such results are also in line with previous studies reported increased functional connectivity between TPJ and near parietal brain areas in functional connectivity analyses (Decety et al., 2008b; Igelstrom and Graziano, 2017).

### FC on TPJ allows to classify OT and PL group

SVM classification with the resting-state FCs has been largely used for detecting brain diseases (Zeng et al., 2012), predicting brain maturity (Dosenbach et al., 2010), or mental states (Tagliazucchi and Laufs, 2014). Here we adopt it to investigate whether the TPJ related functional connectivity allows to robustly differentiate OT and PL group at the individual level. Impressively, using only eight most discriminative FCs, rather than a large number of FC features, can successfully detect 74.58% of participants. It confirms that these FC changes are caused by OT effect, and the effects are reliable at individual level. FC between lTPJ and rRect has highest contribution to the classification, as the connection is higher in PL group than OT group. Existing network efficiency analysis finding showed decreased connection strength between right temporal, parietal and occipital lobes in high-risk of autism spectrum disorder (ASD), without report stronger connection in ASD (Lewis et al., 2014). Combining the findings of increased lTPJ-lIPL, rTPJ-lPrec, rTPJ-rSPL, rTPJ-rPrec, our finding may indicate a OT effect of strengthen ipsilateral brain connection, especially from the right TPJ, while weaken contralateral brain connection from left TPJ.

### Empathic concern intermediates OT effects on FC

Our results show that drug has an interaction effect on the FC between TPJ and Rectus (see Figure 4). Specifically, lTPJ-RECT resting state connectivity positively correlated with empathic concern in PL group. Using task related fMRI, previous study has shown that brain activity in ventral MPFC, including gyrus rectus, positively correlated with empathic ability (Schulte-Ruther et al., 2011). Also, it has been reported that TPJ activity positively correlated with the ingroup empathy bias (Cheon et al., 2011). Considering these findings are based on OT–free subjects, they explain the positive correlation between TPJ-rectus FC and empathic concern in PL group but not in OT group.

### Limitation and future work

Potential limitation and future work should be recognized. A large limit of this study is that we did not collect resting fMRI data before and after the OT administration, thus this study cannot directly show a causal effect of OT on functional connectivity. Future work with the comparisons from the same group of subjects should be conducted. Moreover, ICA cleanup in data preprocessing, such as FIX implemented in FSL (Salimi-Khorshidi et al., 2014), can be applied in the future to remove noise effects before FC analyses. Another limitation of this work is that the input features for SVM are identified from FC images, and we did not perform any type of feature selection during training procedure. Filter methods or wrapper methods to select optimal FC features (Kassraian-Fard et al., 2016) from a large FC pool might be potential way to further improve the accuracy of classification.

## ACKNOWLEDGEMENT

This work was supported by the National Natural Science Foundation of China [grant number: U1736125, 31400963], and CAS Key Laboratory of Behavioral Science, Institute of Psychology to HW. QL was supported by James Boswell fellowship and FWO fellowship.

---

**Supplementary Figure 1.**
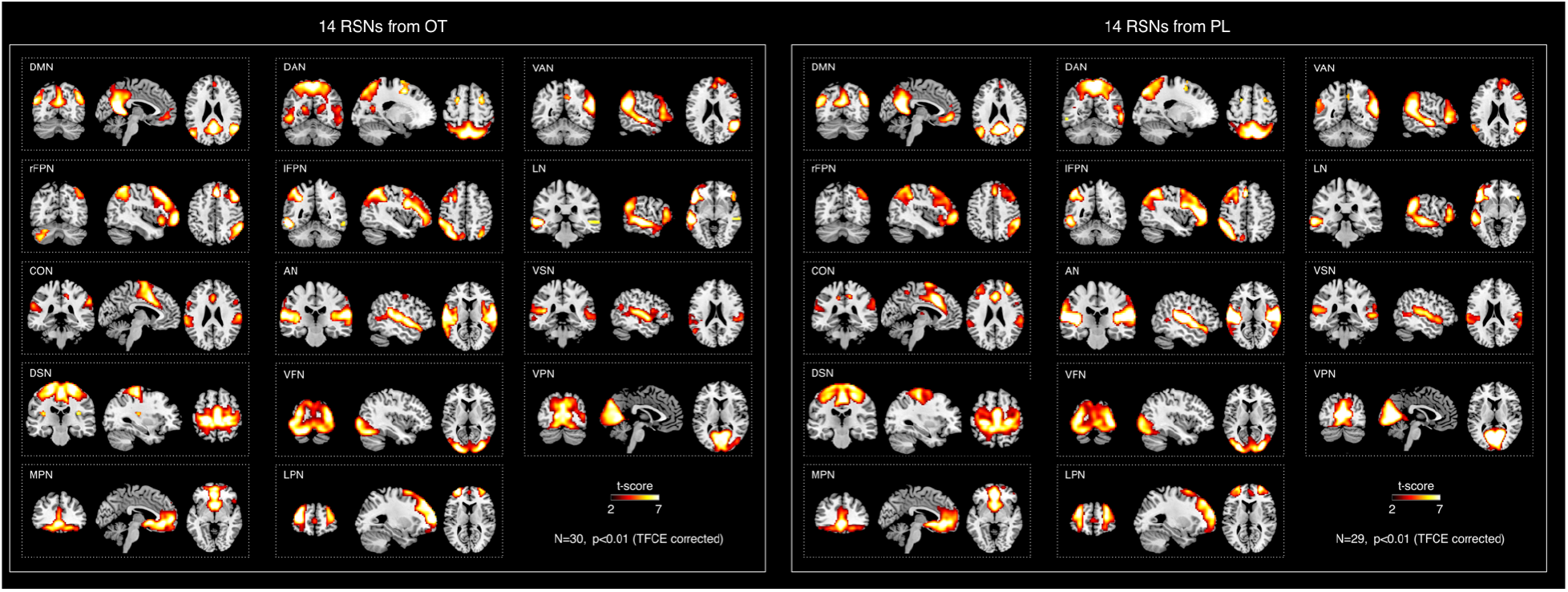
14 RSNs extracted from OT group and PL group. The RSN maps were selected from IC maps which extracted from ICA analysis on BOLD signal time series for each individual, by a template-matching strategy. The group-level RSN maps from OT group and PL group were presented. Surprisingly, the spatial maps showed no topological differences and no brain regions survived from TFCE correction, which implies that OT has no effects on the topology of large-scale RSNs.

**Supplementary Figure 2.**
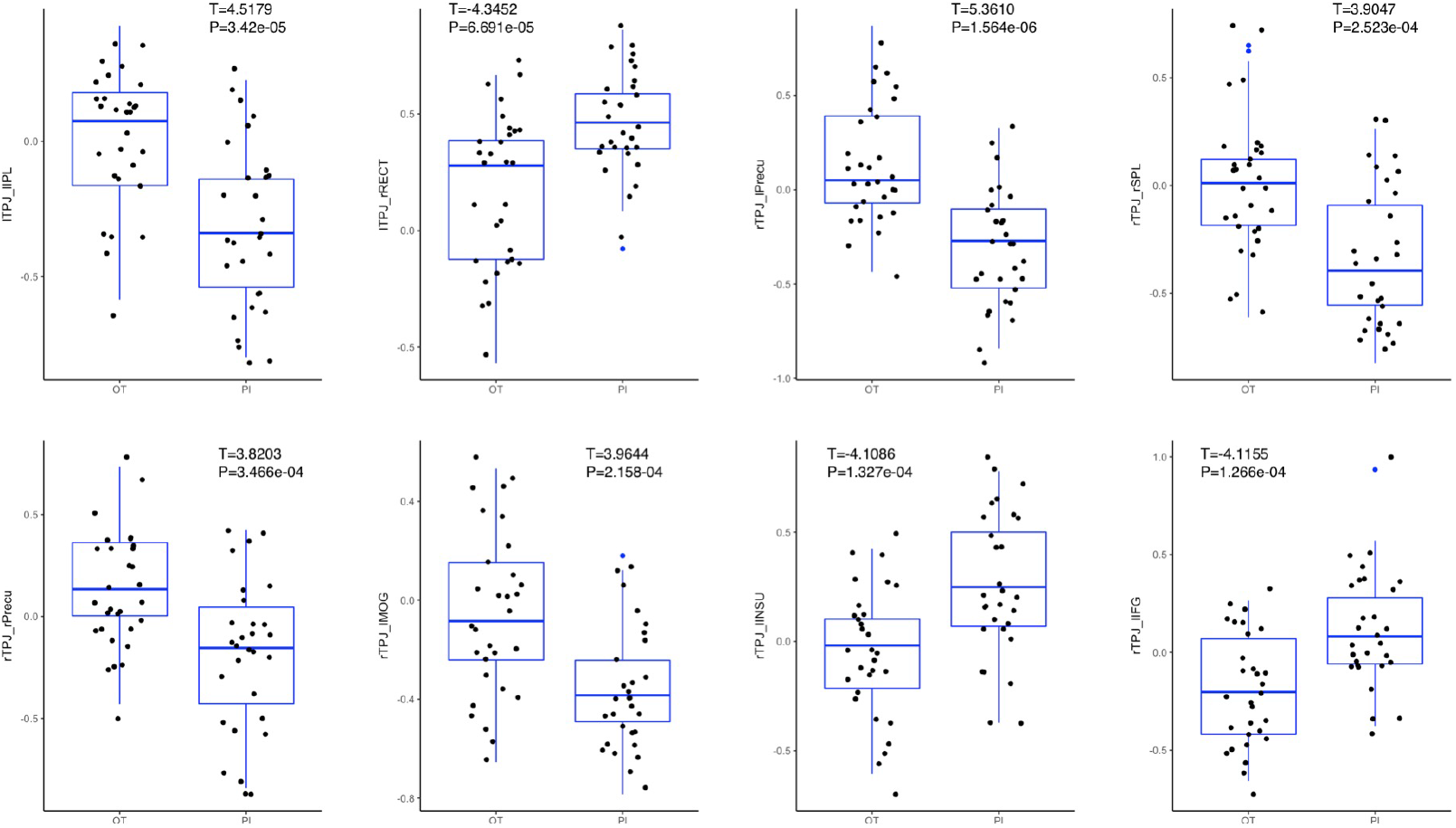
Boxplot of functional connectivity between the identified eight pairs of regions.

**Supplementary Figure 3.**
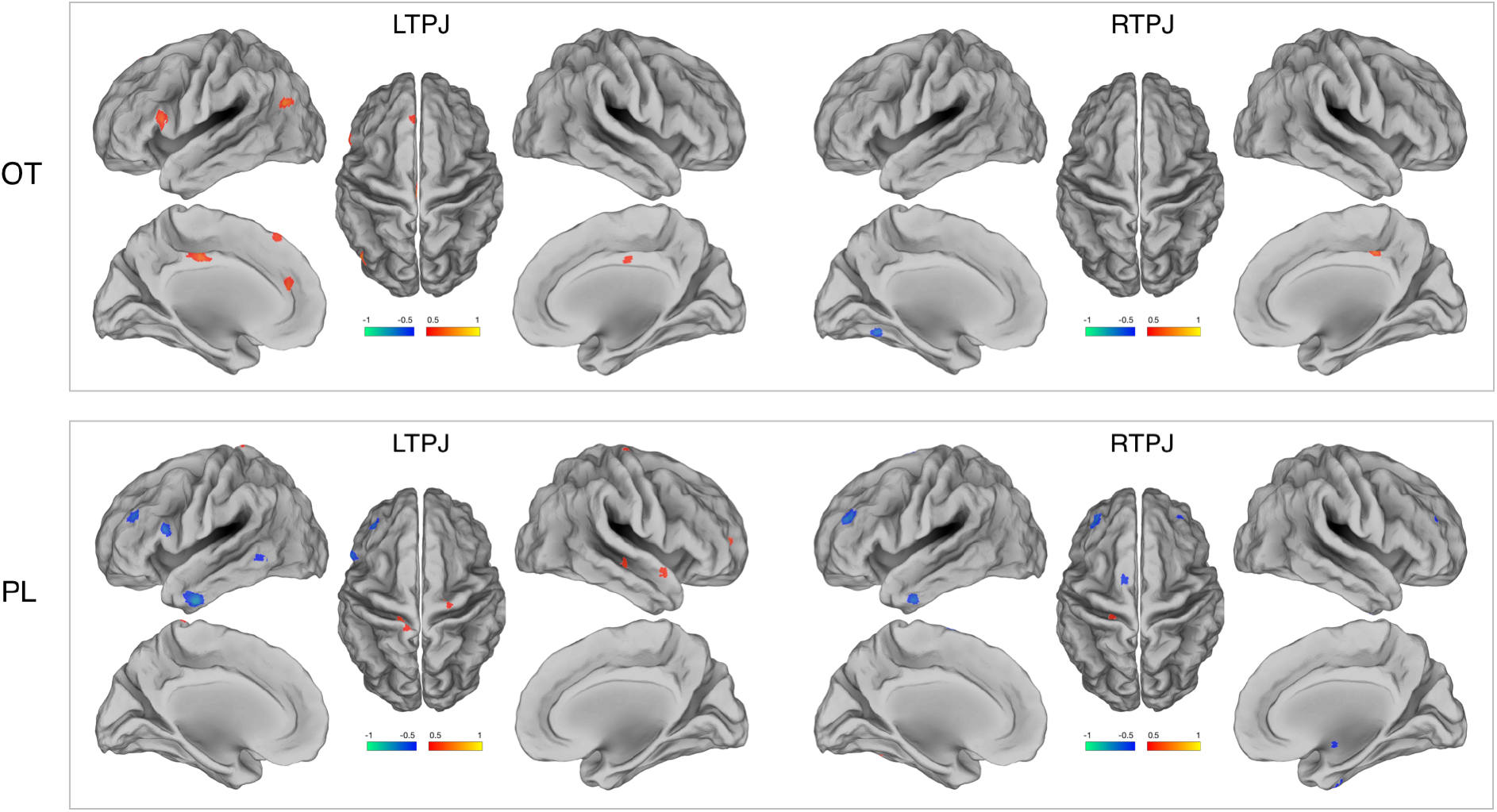
The relationship between perspective taking and the TPJ connectivity. The brain regions correlated (threshold at r > 0.5, labeled with yellow/red) or anitcorrelated (threshold at r < -0.5, labeled with green/blue) with perspective taking scale are indicated on the map.

**Supplementary Figure 4.**
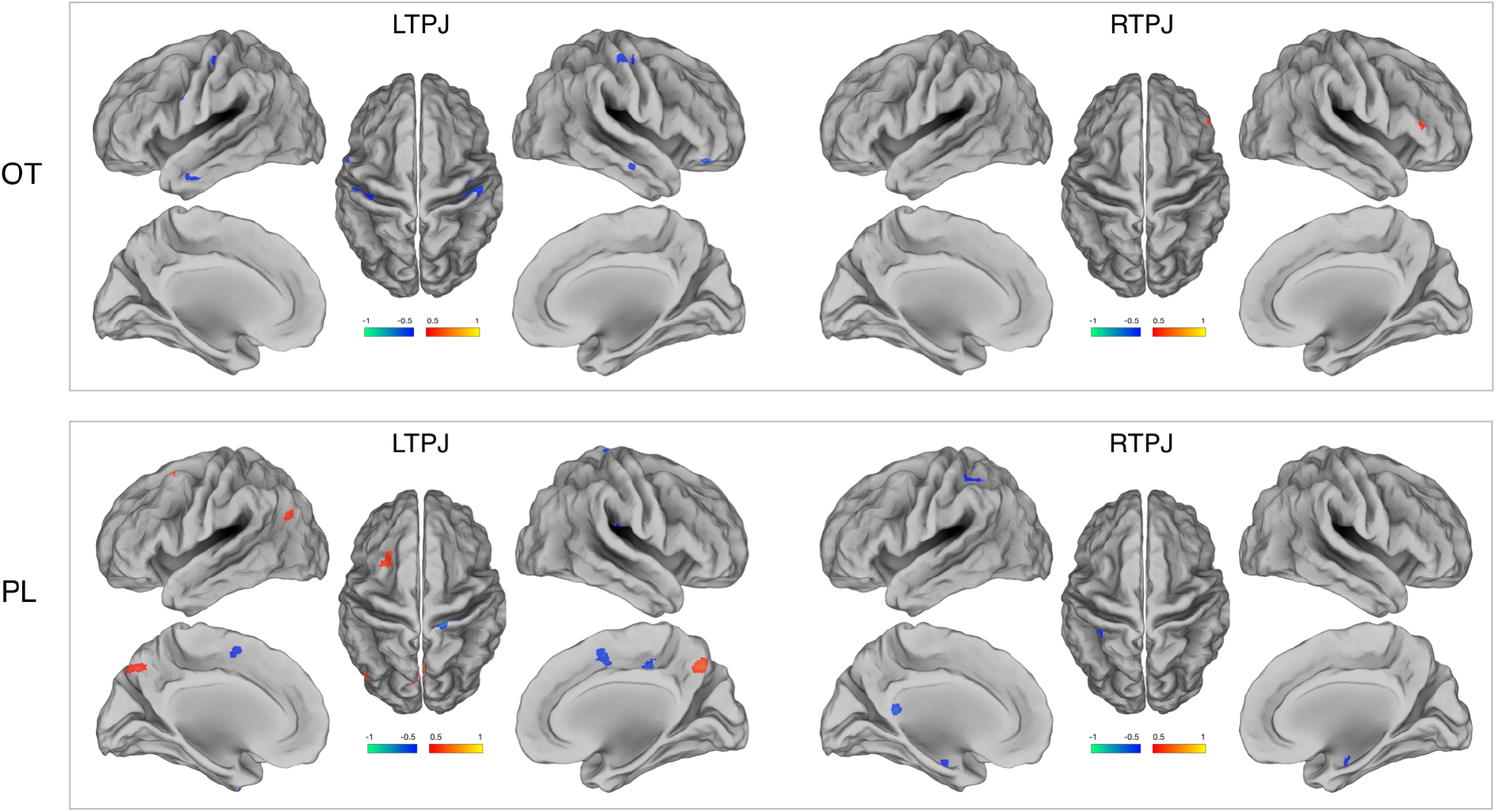
The relationship between fantasy scale and TPJ connectivity. The brain regions correlated (threshold at r > 0.5, labeled with yellow/red) or anitcorrelated (threshold at r < -0.5, labeled with green/blue) with fantasy scale are indicated on the map.

**Supplementary Figure 5.**
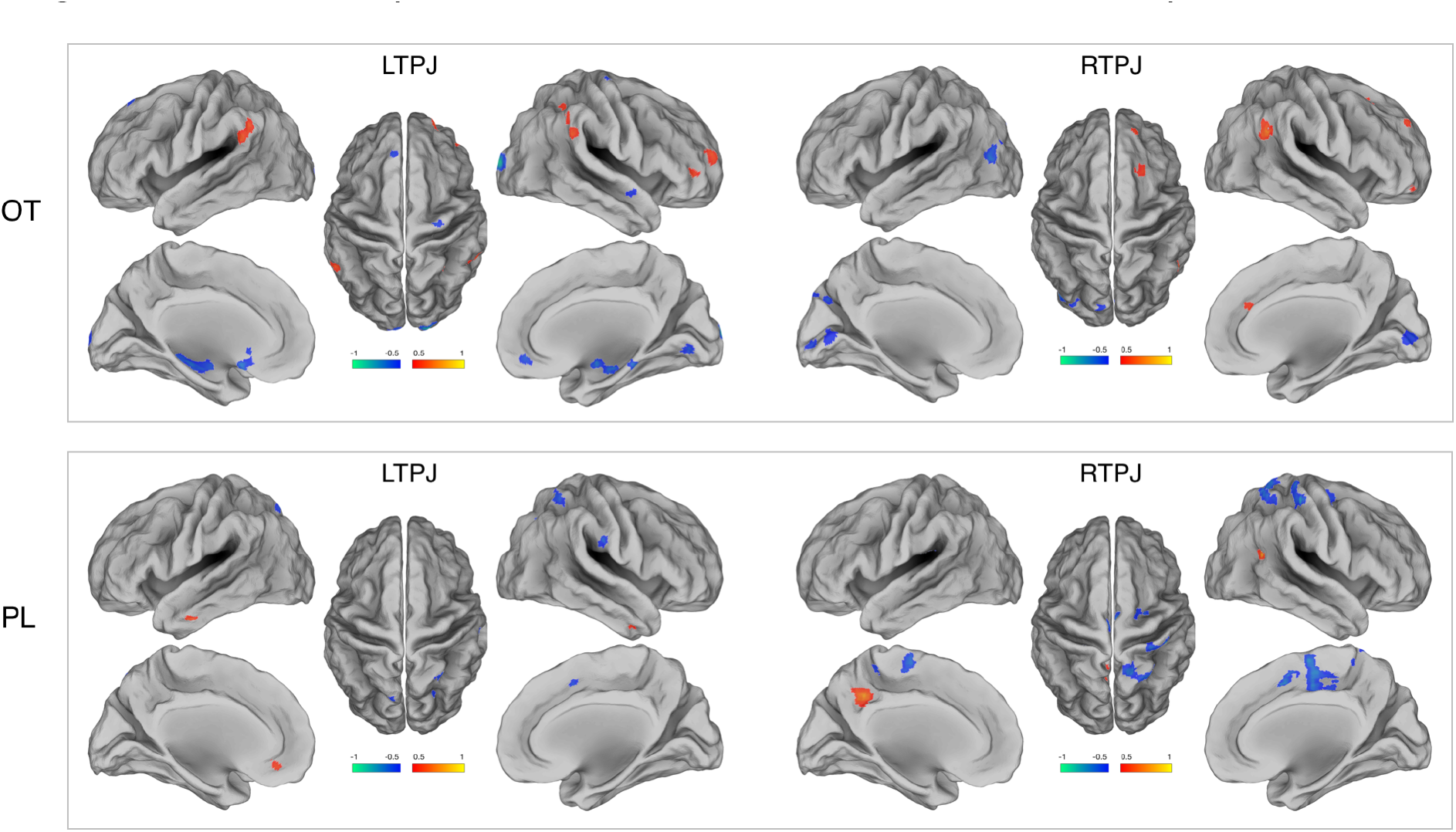
Empathic concern mediates the TPJ connectivity. The brain regions correlated (r > 0.5, labeled with yellow/red) or anitcorrelated (r < -0.5, labeled with green/blue) with empathic concern scale are indicated on the map.

**Supplementary Figure 6.**
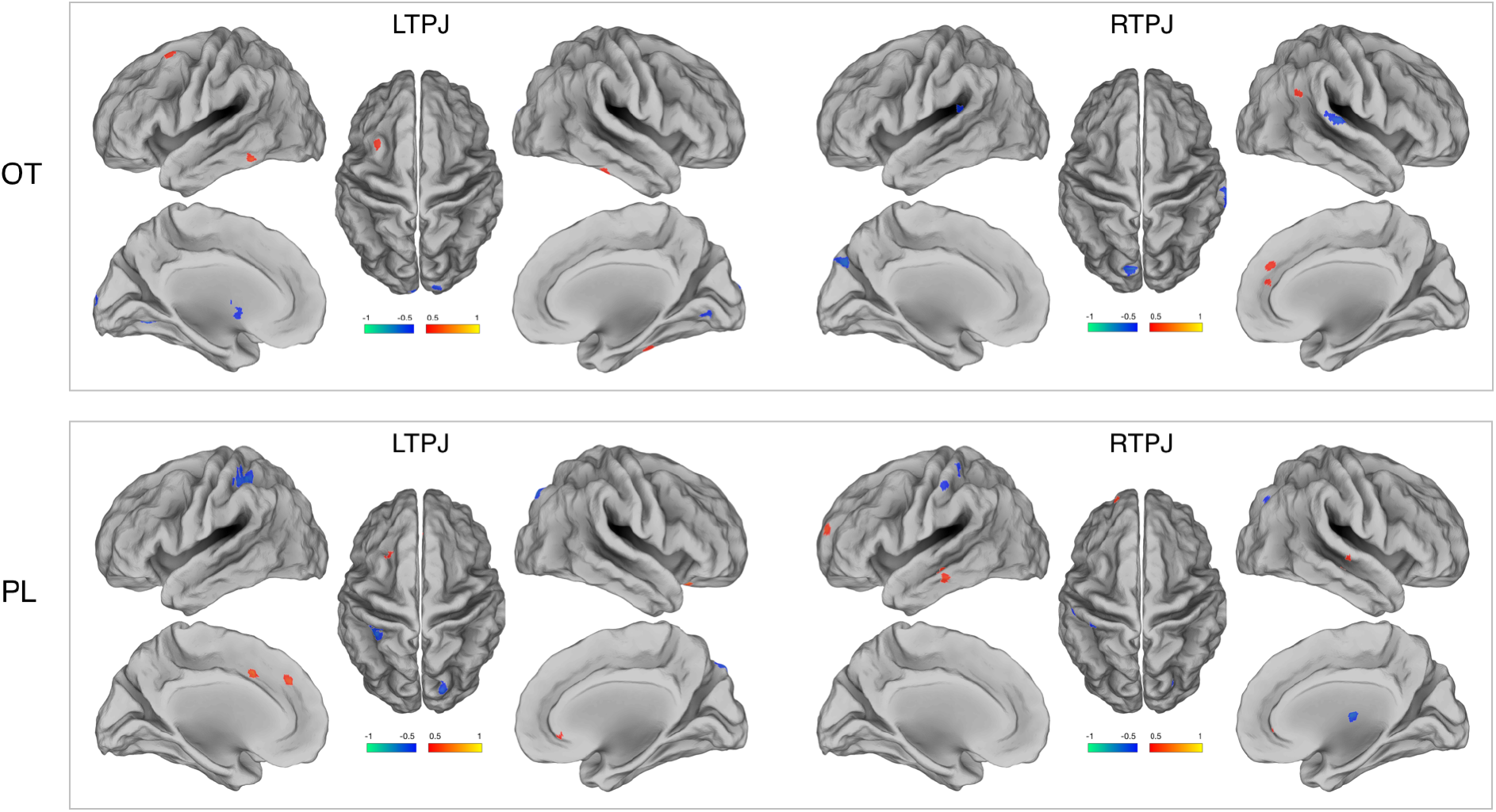
Personal distress mediates the TPJ connectivity. The brain regions correlated (r > 0.5, labeled with yellow/red) or anitcorrelated (r < -0.5, labeled with green/blue) with personal distress scale are indicated on the map.

